# Integrated genomics and comprehensive validation reveal novel drivers of genomic evolution in esophageal adenocarcinoma

**DOI:** 10.1101/2020.04.23.058230

**Authors:** Subodh Kumar, Leutz Buon, Srikanth Talluri, Marco Roncador, Chengcheng Liao, Jiangning Zhao, Jialan Shi, Chandraditya Chakraborty, Gabriel B. Gonzalez, Yu-Tzu Tai, Rao Prabhala, Mehmet K. Samur, Nikhil C. Munshi, Masood A. Shammas

## Abstract

Identification of genes driving genomic evolution can provide novel targets for cancer treatment and prevention. Here we show identification of a genomic instability gene signature, using an integrated genomics approach. Elevated expression of this signature correlated with poor survival in esophageal adenocarcinoma (EAC) as well as three other human cancers. Knockout and overexpression screens confirmed the relevance of this signature to genomic instability. Indepth evaluation of TTK (a kinase), TPX2 (spindle assembly factor) and RAD54B (recombination protein) further confirmed their role in genomic instability and tumor growth. Mutational signatures identified by whole genome sequencing and functional studies demonstrated that DNA damage and homologous recombination were common mechanisms of genomic instability induced by these genes. Consistently, a TTK inhibitor impaired EAC cell growth in vivo, and increased chemotherapy-induced cytotoxicity while inhibiting genomic instability in surviving cells. Thus inhibitors of TTK and other genes identified in this study have potential to inhibit/delay genomic evolution and tumor growth. Such inhibitors also have potential to increase chemotherapy-induced cytotoxicity while reducing its harmful genomic impact in EAC and possibly other cancers.

---

Esophageal adenocarcinoma (EAC) is the sixth biggest cause of cancer deaths throughout the world^1^. The incidence of this cancer is rising rapidly in Western countries. Moreover, the tools and strategies for early detection and treatment of this cancer are still not very effective and disease remains to have poor clinical outcome^1^. The disease is associated with a precancerous condition, Barrett’s esophagus (BE), which gradually progresses to EAC^2^. Like most cancers, EAC is also associated with a marked genomic instability which arises at an early stage and allows ongoing accumulation of genomic changes, some of which contribute to development and progression of cancer^3^. Evaluation of single nucleotide polymorphisms in patient genomes has shown that in addition to EAC, genomic changes can also be seen in majority of BE cases^4^. Microsatellite instability has also been detected at both the BE as well as EAC stages^5^. In fact changes of both the genetic and epigenetic type have been detected at BE stage^6^. There is also evidence that genomic instability increases with progression of BE to cancer. Consistently, the aneuploidy detected in a subset of BE cases, has been shown to progressively increase with progression to EAC^7^. Similarly, it has been reported that copy number changes spanning relatively large areas of genome are rare in early stage disease but occur more frequently and involve larger areas of genome in advanced stages^8^. Moreover, it has been shown that although mutational load in precancerous BE cases is lower than EAC, it is higher than that observed in certain other cancer types^9,10^.

EAC genome is highly aberrant with ∼ ten single-nucleotide variations per million base pair^9^. Genomic instability and its consequences can probably be attributed to chemoresistant nature of EAC^11^. There is also evidence which suggests that genomic instability contributes to disease progression^12^ and associates with poor survival in EAC^13^. Using EAC and multiple myeloma as cancer model systems, we have reported that homologous recombination (HR), a prominent DNA repair system, is spontaneously elevated in these cancers and contributes to genomic instability^14-16^, development of drug resistance^16^, and telomere length maintenance and tumor growth^17^. We have also shown that acid and bile, the main contents of gastroesophageal refluxate, induce HR activity in human cells^14^. Bile acids have also been shown to cause oxidative DNA damage in BE cells^18^. Since DNA damage can activate HR, specific microenvironment of BE/EAC (exposed to acid and bile) may contribute to dysregulation of HR and genome stability. In this study, we used an integrated genomics approach to identify novel mediators of genomic instability in EAC. Functional significance of these genes was confirmed in knockout and overexpression screens. Three of these genes (TPX2, TTK and RAD54B) representing diverse pathways were further evaluated in vitro and in vivo. We demonstrate that novel genes identified in this study and their inhibitors (such as TTK inhibitor used in this study) have potential to inhibit/reduce genomic instability and growth of cancer cells in vitro and in vivo. Such inhibitors also have potential to reduce chemotherapy-induced genomic instability, while increasing their cytotoxicity.

## Results

### Integrated genomics identifies a genomic instability gene signature

The stepwise process involved in the identification/validation of genes is presented in **Figure 1 (I-II)**. Briefly, the integration of copy number, expression and survival data identified 31 genes which were overexpressed in EAC, and whose elevated expression correlated with increased genomic instability (as assessed from total copy number events) in each patient. Figure 1B shows that elevated expression of 31 gene signature (GIS31) is associated with increased risk in TCGA patient dataset. Pairwise correlation also identified several subgroups of these genes showing co-overexpression. Two such examples (CDK1, TROAP, KIF4A, STIP1 and ERCC6L, MST4, NEK2) are shown by red rectangles in **Supplementary Figure 1**. Elevated expression of this gene signature also correlated with poor overall survival in pancreatic cancer, lung cancer, three different clinical datasets of multiple myeloma as well as in a second EAC patient dataset (GSE19417; Figure 1C, I-VI). Top pathways related to these genes were cell cycle, cell division, ATP binding and organelle (including chromatin and spindle) organization (Figure 1D), indicating the importance of these pathways with relevance to genomic evolution.

**Figure 1.**
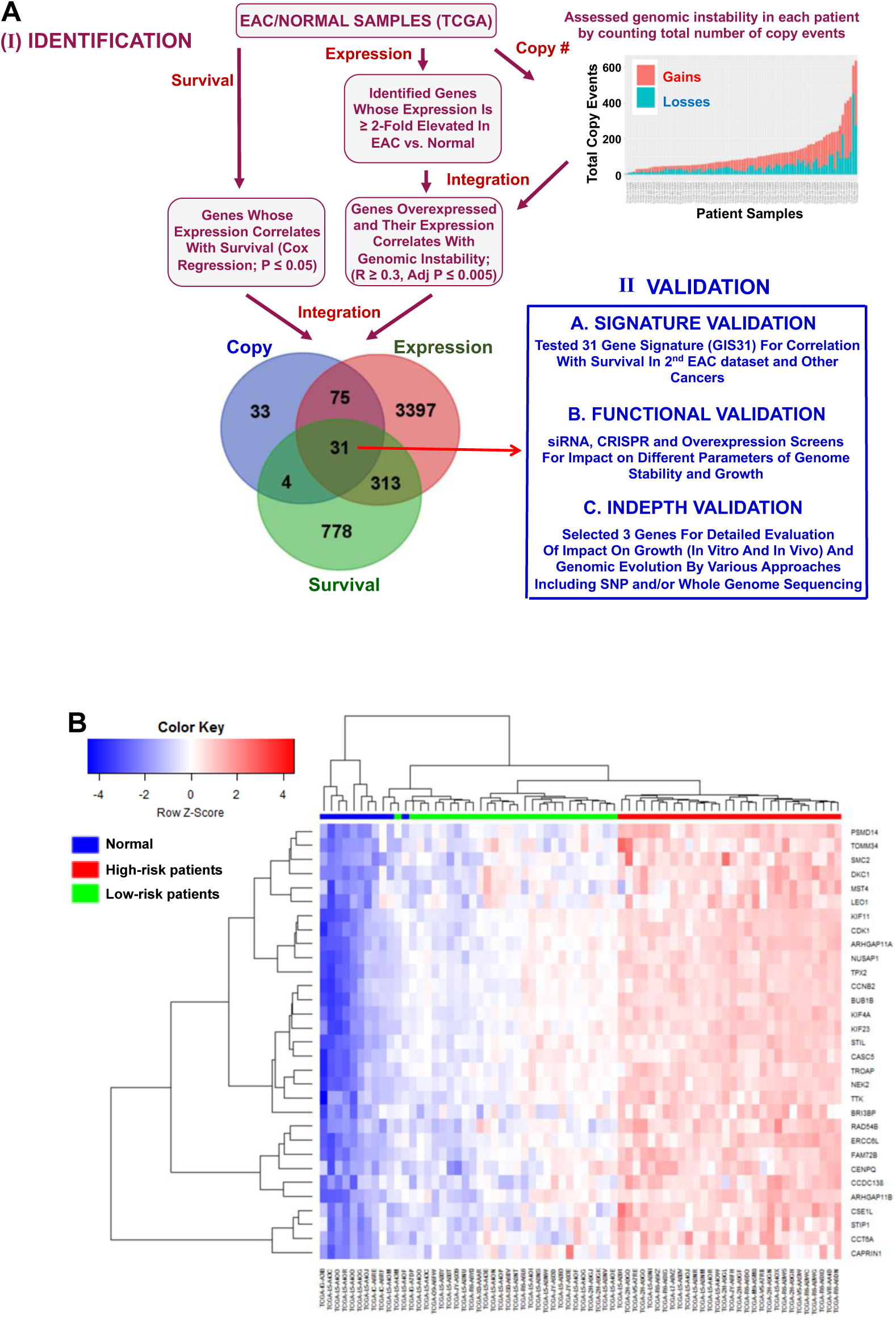
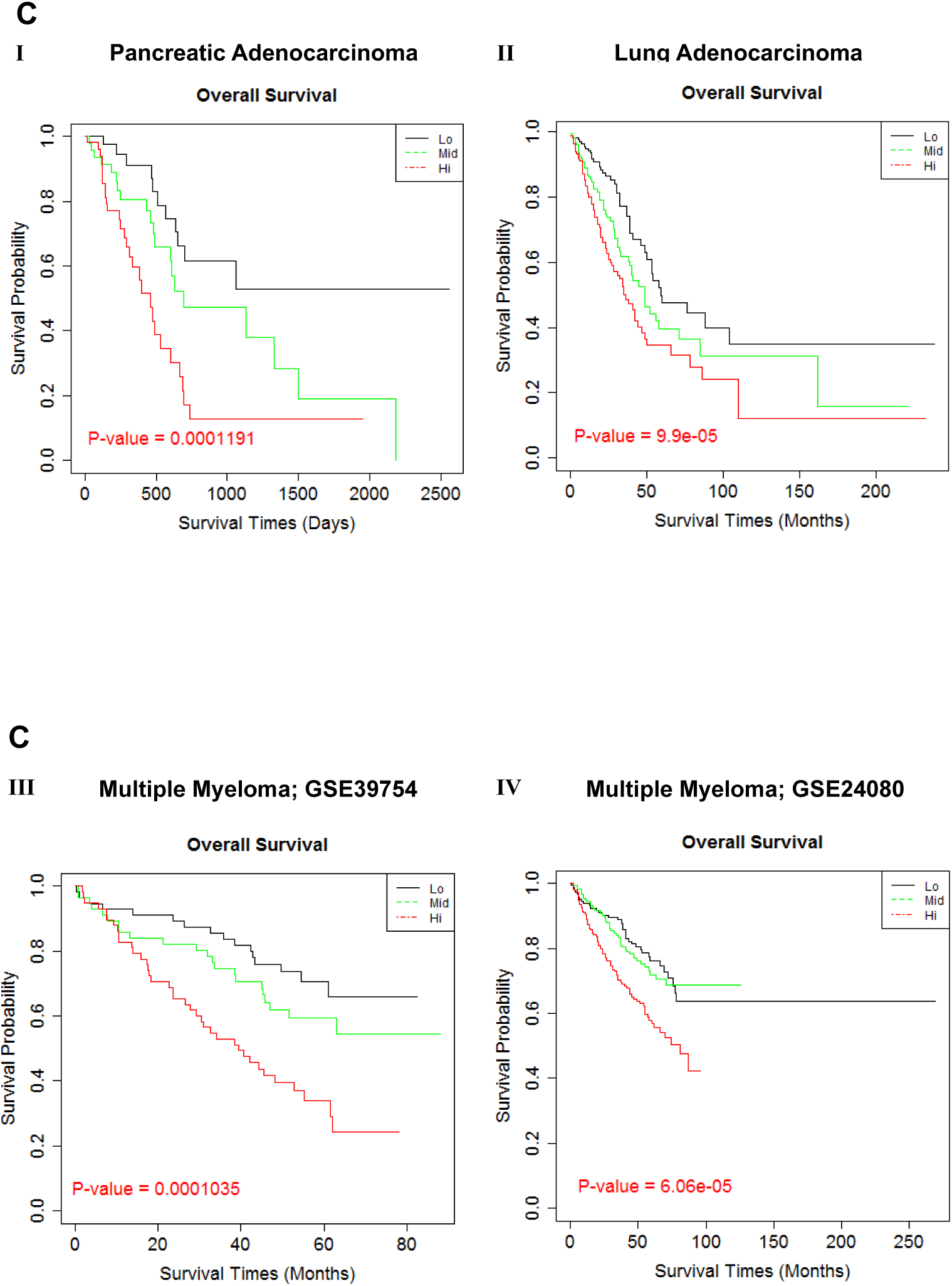
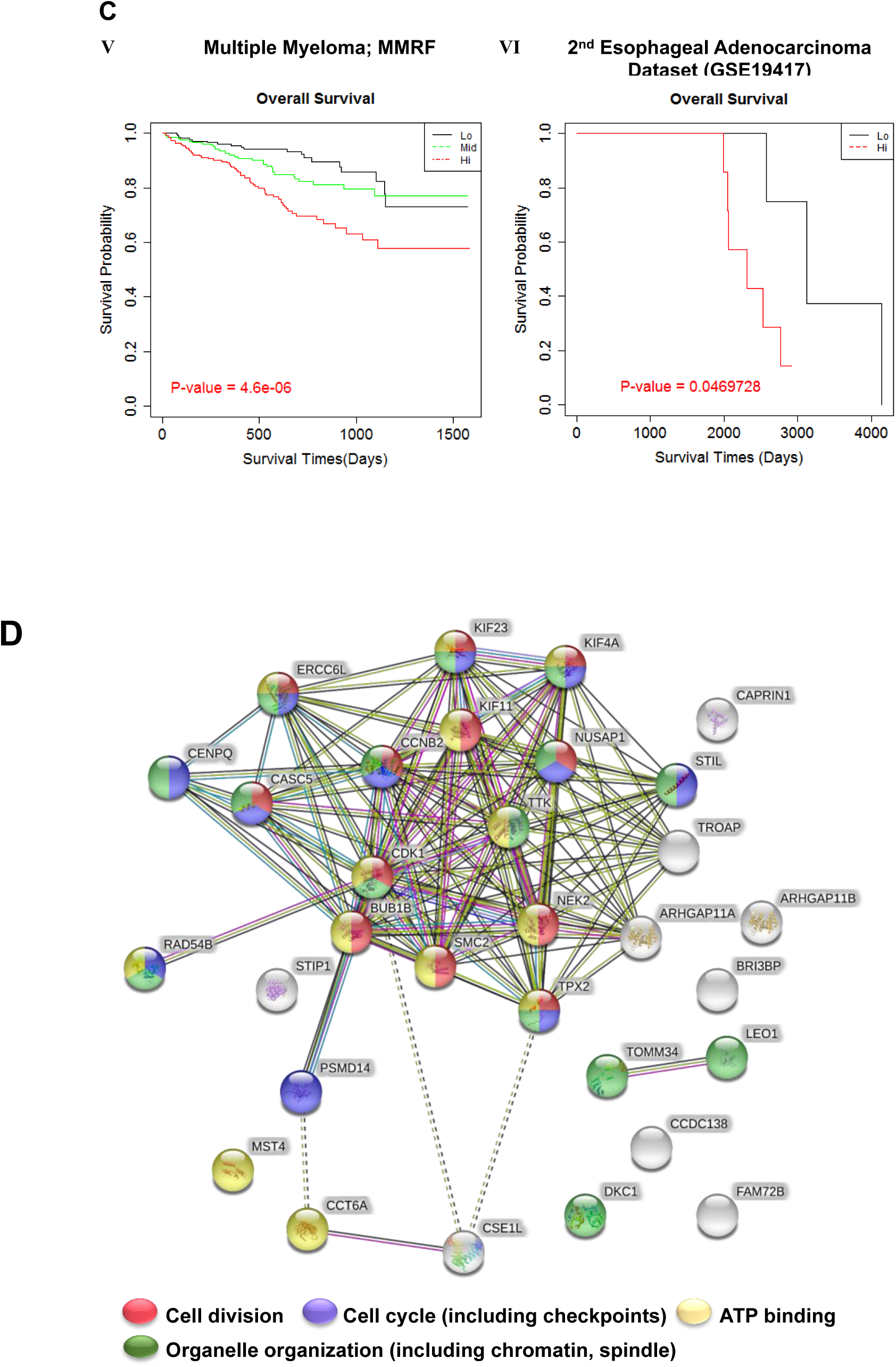
Identification of novel drivers of genomic evolution in EAC. **(A) Experimental design. Panel I:** Stepwise process involved in the identification of genes. **1)** Assessed gene expression in 11 normal and 88 EAC patient samples in TCGA dataset and identified genes that were overexpressed in EAC; **2)** Assessed genomic instability in patient samples by counting total number of copy events in each patient. Integrated genomic instability data with expression data to identify genes that were overexpressed in EAC and whose expression correlated with genomic instability; **3)** Evaluated resulting genes for correlation with survival. This led to identification of 31 genes that were overexpressed in EAC and whose expression correlated with genomic instability and overall survival in EAC patients in TCGA dataset; **Panel II:** Describes plan for functional validation of genes; 1) The expression of 31 gene signature was tested for correlation with survival in a second EAC dataset as well as in other cancers; 2) Conducted siRNA, CRISPR/Cas9 and overexpression screens for impact on different parameters of genome stability and growth; 3) Selected 3 genes for detailed evaluation of impact on genomic evolution by various approaches including SNP and/or whole genome sequencing; 4) Evaluation in SCID mice. **(B)** Heat map shows that expression of 31 genes is high in EAC patients with poor overall survival (red bar) and low in patients with relatively better survival (green bar) (in TCGA dataset) and also low in normal samples (blue bar). **(C)** Elevated expression of thirty one gene signature (identified in TCGA patient dataset) correlated with poor overall survival in pancreatic cancer (I), lung cancer (II), three different clinical datasets of multiple myeloma (III-V) as well as in a second EAC dataset (GSE19417). **D)** Interconnectivity among GIS31 genes (based on STRING networks) is shown by lines and shared functional roles indicated by different colors.

### Functional screens confirm relevance of GIS31 genes with genomic instability

Using EAC and multiple myeloma as model systems we have demonstrated that spontaneously elevated homologous recombination (HR) contributes to genomic instability^14^, telomere length maintenance and tumor growth^17^, and development of drug resistance^16^. To investigate the functional relevance of GIS31 genes, we first conducted an siRNA screen (using validated siRNAs from Sigma) to evaluate their role in HR activity in EAC cells. Suppression of 19 out of 31 (61%) genes resulted in significant inhibition of HR activity (p < 0.05) (**Supplementary Figure 2**). Fourteen of these genes were further evaluated in loss of function (using CRISPR/Cas9 system) and/or gain of function screens for their role in HR, micronuclei (a marker of ongoing genomic rearrangements and instability^19^) and cell viability. For CRISPR/Cas9 loss of function screen, FLO1 cells stably integrated with Cas9 were transduced with guide RNAs (2 – 3 per gene) and impact of gene-knockdown on HR and cell viability investigated. Relative to average of two control guides, transduction with each guide RNA caused significant inhibition of HR activity (ranging from 27 − 71% inhibition; p < 0.03) (Figure 2A, I), in three independent experiments. Although BUB1B did not show any impact in siRNA screen, its knockdown by CRISPR/Cas9 resulted in significant inhibition of HR activity by all three guides. For all other genes, the knockdown by CRISPR/Cas9 or siRNA showed similar impact of these genes on HR, establishing their functional significance in genome maintenance. Except NEK2, CENPQ and NUSAP1, suppression of all other genes was also associated with significant inhibition (P < 0.05) of cell viability (Figure 2A, II). Eleven genes were overexpressed in FLO1 cells and impact on HR activity and micronuclei (a marker of genomic instability) evaluated. Overexpression of each of these genes resulted in significant increase (ranging from 29% to 90% increase; P ≤ 0.04) in HR activity (Figure 2B, I). Moreover, the increase in HR following overexpression correlated with its inhibition following knockdown of the same set of genes (R^2^ = 0.7), further supporting their role in HR (Figure 2B, II). Overexpression of these genes was also evaluated for impact on micronuclei (another marker of genomic instability). Panel I in Figure 2C shows representative images of nuclei and micronuclei in FLO1 cells overexpressing GIS31 genes, whereas bar graph (in Panel II) summarizes the results from three independent experiments. Overexpression of six out of eleven genes tested was associated with significant (1.8 − 3.5-fold; p < 0.04) increase in micronuclei (Figure 2C, II). Overall 65% of identified genes had a role in one or more activities evaluated, confirming the functional relevance of identified signature to genome stability and growth.

**Figure 2.**
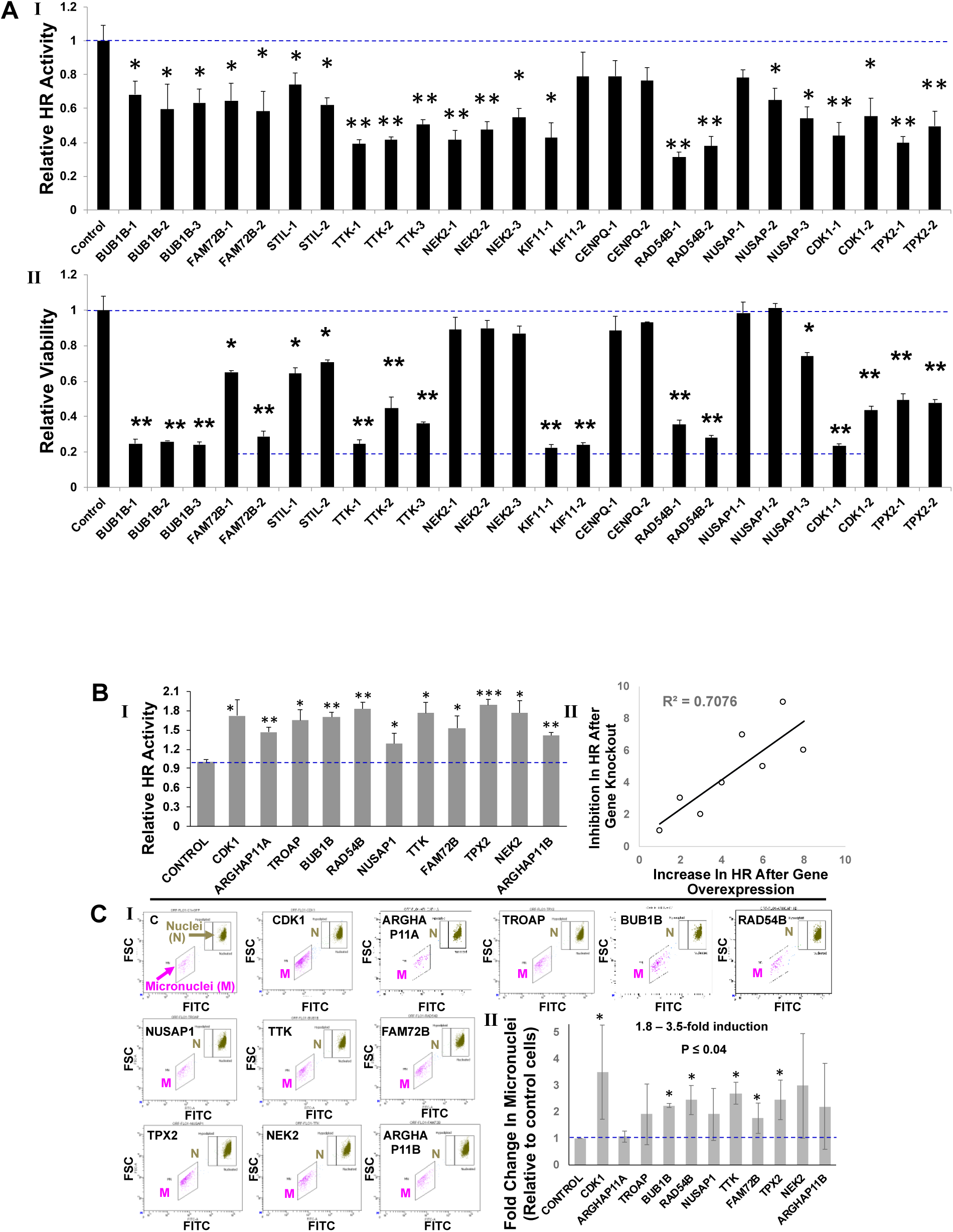
Knockdown and overexpression screens evaluating GIS31 genes for impact on different aspects of genome stability and cell viability. **(A) <u>Knockdown</u>**: EAC (FLO1) cells, stably transduced with Cas9, were treated with lentiviral single-guide RNAs, either non-targeting control or those targeting specific genes (two or three guides per gene). Following selection, cells were cultured for 10 days and evaluated for impact on HR activity (I) and cell viability (II), relative to control (control bar represents average of two different control guides); Error bars represent SDs of three experiments. Two-tailed p values (*, p < 0.05 – 0.001; **, p < 0.001) indicate significance of difference relative to control. **(B-C) <u>Overexpression</u>**: FLO1 cells were treated with lentiviral constructs, either control or those carrying open reading frames (for overexpression) of genes, selected in puromycin, and evaluated for HR (B) and micronuclei (a marker of genomic instability) (C). **(B) I**. Bar graph showing HR activity, relative to control; Error bars represent SDs of three independent experiments. Two-tailed p values: *, p < 0.05; **, p < 0.0007; ***, p < 0.00005; **II**. The line plot shows correlation of increase in HR following overexpression vs. its inhibition following knockdown of the same set of genes (shown in panel A). Note: For knockdown, the average inhibition of HR activity by all (2 or 3) guides for each gene was used. **(C) I**. Representative images showing nuclei (N) and micronuclei (MN; a marker of genomic instability) in transduced cells. X-axis is FITC signal indicating amount of DNA in nuclei or micronuclei and Y-axis (FSC or forward scatter) indicates size which distinguishes nuclei from micronuclei; **II**. Bar graph showing fold change in micronuclei, relative to control cells; Error bars represent SDs of three experiments. * = two-tailed p value < 0.04.

Of genes with significant impact on these activities, TTK (a dual specificity protein kinase), TPX2 (a spindle assembly factor required for normal assembly of mitotic spindles and microtubules) and RAD54B (a recombination protein), representing diverse pathways, were selected for further evaluation as described below.

### TTK, TPX2 and RAD54B are overexpressed in EAC cell lines and patient samples

Expression of TTK, TPX2 and RAD54B, as evaluated by real time PCR, is elevated >2-fold (P<0.05) in all three EAC cell lines, relative to normal primary human esophageal epithelial cells (**Figure 3A, I**). Error bars indicate SDs of experiments conducted in triplicate. The expression of all three genes is also significantly elevated in EAC patients, relative to normal samples (P<0.00005; Figure 3A, II).

**Figure 3.**
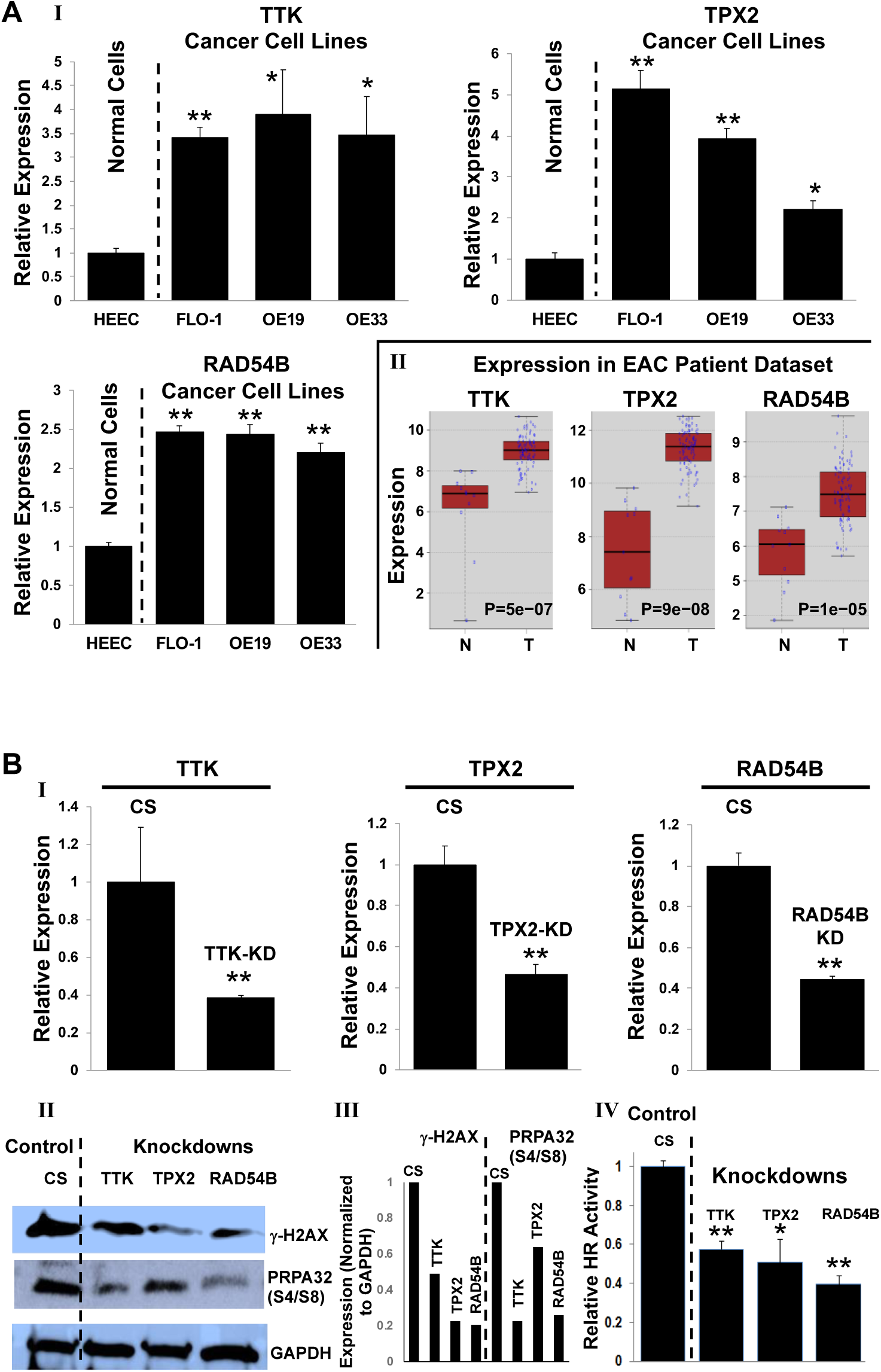
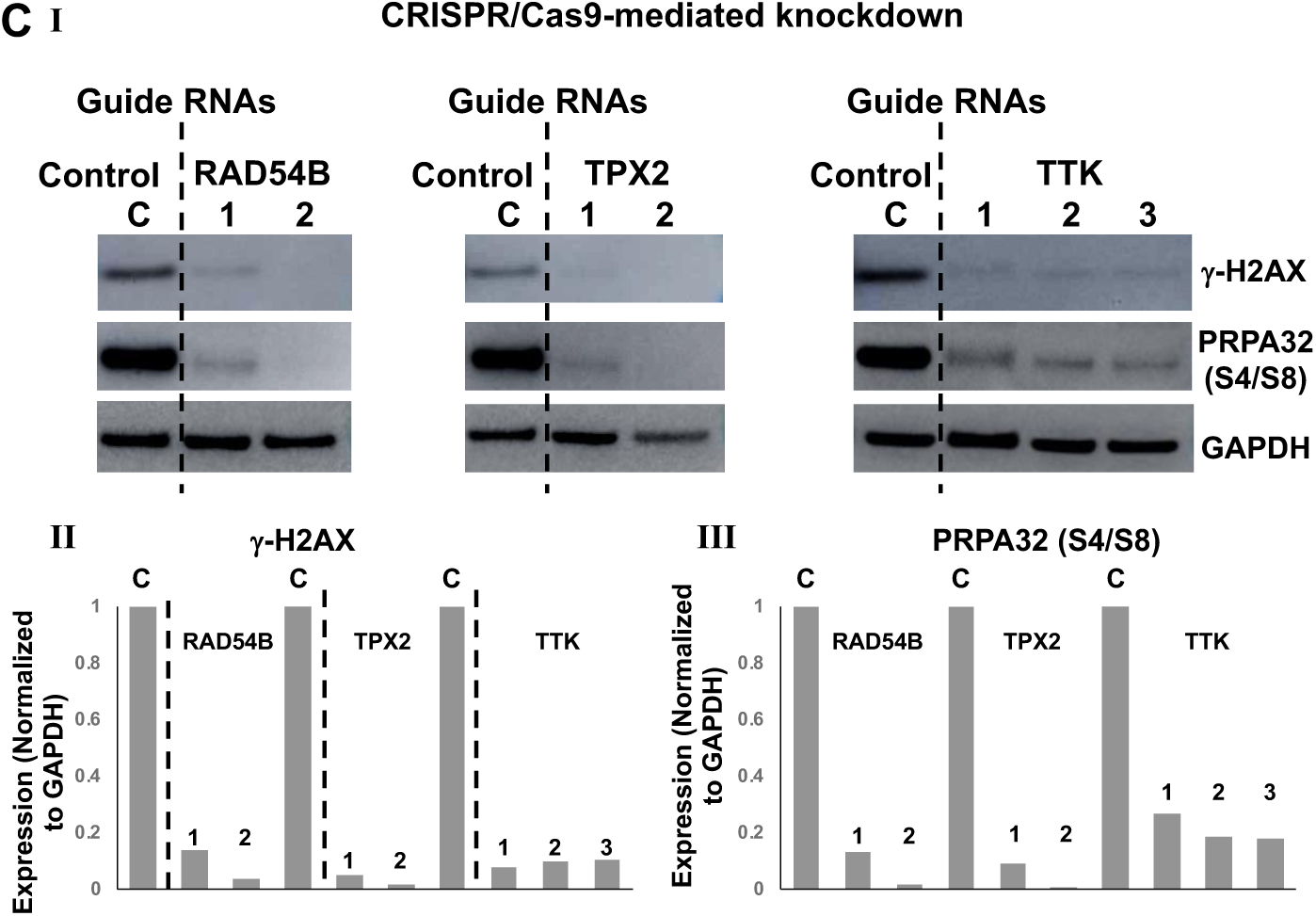
Suppression of elevated TTK, TPX2 or RAD54B expression in EAC cells inhibits spontaneous DNA breaks, HR and growth in vitro and in vivo. **(A) I**. Gene expression, evaluated in normal primary human esophageal epithelial (HEsEpi) cell and EAC cell lines (FLO-1, OE19 and OE33), using real time PCR; Error bars indicate SDs of experiments conducted in triplicate. Two-tailed p values: * = p < 0.05 - 0.001; ** = p < 0.001. **II**. Log2 gene expression in EAC relative to average expression in normal samples in TCGA patient dataset (n=88). **(B) shRNA-mediated suppression of TTK, TPX2 or RAD54B reduces spontaneous DNA breaks and HR in EAC cells**. FLO-1 cells were transduced with lentivirus-based shRNAs, either control (CS) or those targeting TTK, TPX2 or RAD54B, and following selection, knockdown (KD) of gene expression confirmed by RT PCR (I), levels of **γ**-H2AX and phosphorylated RPA32 evaluated by Western blotting (II-III), and HR activity measured using a functional assay (IV). Error bars in panel I indicate SDs of experiment conducted in triplicate and those in panel IV are SDs of three experiments. Two-tailed p values: * = p < 0.05 - 0.001; ** = p < 0.001. **Panels (II-III)**. Western blot image (II) and quantitation of the blot (III) are shown. **(C) CRISPR/Cas9-mediated suppression of TTK, TPX2 or RAD54B reduces spontaneous DNA breaks and DNA end resection in EAC cells**. FLO1 cells, stably transduced with Cas9, were treated with lentiviral single-guide RNAs, either non-targeting control (C) or those targeting these genes (two or three guides per gene). Following selection in puromycin, cells were cultured for 10 days and impact of their suppression on levels of **γ**-H2AX (a marker of DNA breaks) and pRPA32 (a marker of DNA end resection) evaluated by Western blotting. Images (I) and the bar graphs presenting quantitation of Western blots (II-III) are shown.

### Suppression of TTK, TPX2 and RAD54B inhibits spontaneous DNA damage and HR activity

We suppressed TTK, TPX2 and RAD54B in EAC (FLO-1) cells by shRNAs (**Figure 3B**) as well as CRISPR/Cas9 system (using two guide RNAs per gene; **Figure 3C**). Knockdown of TTK, TPX2 and RAD54B by shRNAs (shown in **Figure 3B, I**) caused the inhibition of **γ**H2AX (a DNA break marker) expression in FLO1 cells by 51%, 77% and 79%, respectively (Figure 3B, II-III). The same blots were also evaluated for phosphorylation (on ser4 and ser8) of RPA32, a marker of end resection^20^, the key step in the initiation of HR. Suppression of all three genes also led to reduction in p-RPA32 expression (by 36 − 77%), indicating inhibition of DNA end resection in these cells (Figure 3B, II-III). Consistently, the HR activity was also significantly reduced following suppression of all three genes (p < 0.05; Figure 3B, IV). Similar observations were made following CRISPR/Cas9-mediated knockdown using two guides/gene (**Figure 3C**). Relative to control guide RNA, each of the guide against all three genes resulted in > 80% reduction in the levels of **γ**H2AX and > 70% reduction in p-RPA32 expression (Figure 3C). Thus suppression of TTK, TPX2 and RAD54B, whether mediated by shRNAs or CRISPR/Cas9 system, confirm their role in increased spontaneous DNA damage and dysregulation of HR in FLO-1 cells.

### Suppression of TTK, TPX2 or RAD54B impairs EAC cell growth *in vitro* as well as *in vivo*

Control and knockdown cells were cultured and cell viability assessed at various intervals. Over a period of 7 days, the suppression of TTK, TPX2 and RAD54B reduced the viability of EAC cells by 54%, 39% and 35%, respectively (**Figure 4A**). To monitor the impact on growth in vivo, knockdown cells were selected in puromycin, cultured for another two weeks and injected subcutaneously in SCID mice and tumor growth measured at indicated intervals. Relative to control cells, the suppression of all three genes (TPX2, TTK and RAD54B) was associated with significant reduction in tumor size (*p* ≤ 0.04) in SCID mice (**Figure 4B-D**).

**Figure 4.**
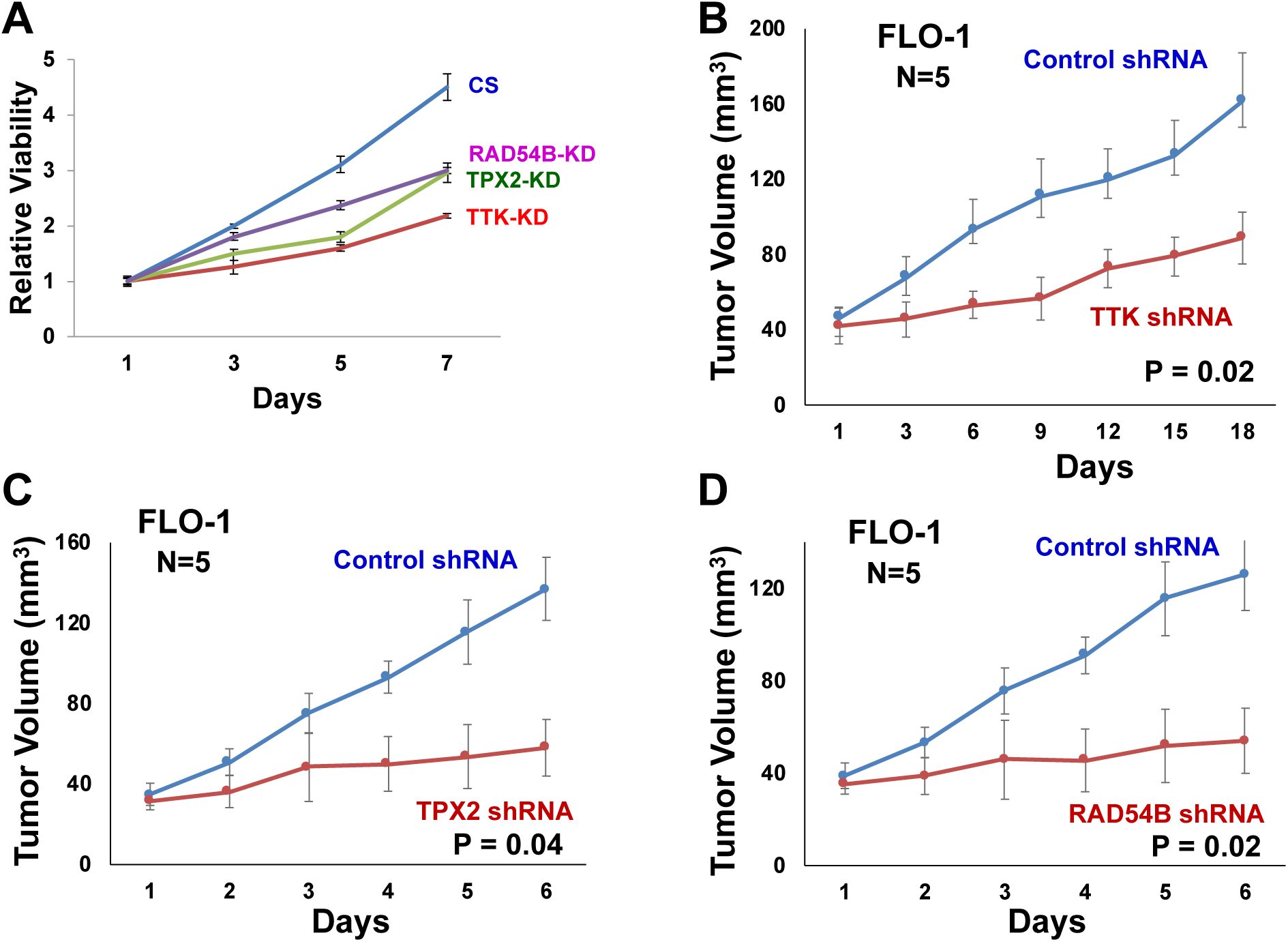
Suppression of TTK, TPX2 or RAD54B impairs EAC cell growth in vitro and in vivo. **(A)** Control and knockdown FLO1 cells were cultured and cell viability assessed at various intervals; **(B-D)** Control and knockdown FLO-1 cells, cultured for > 10 days in vitro, were subcutaneously injected in SCID mice and tumor growth measured at indicated intervals. Error bars indicate SEMs of tumor sizes in different mice. Two-tailed p values ≤ 0.04.

### Transgenic overexpression of TTK, TPX2 and RAD54B in normal esophageal cells increases spontaneous DNA damage and HR activity

Expression of TTK, TPX2 and RAD54B was upregulated in normal primary human esophageal epithelial cells, using lentivirus-based expression plasmids. Overexpression of all three genes (**Figure 5A, I**) led to increase in the expression of **γ**-H2AX (DNA break marker), p-RPA32 (marker of DNA end resection) (Figure 5A, II) as well as significant increase in HR activity (p < 0.001; Figure 5A, III) in normal esophageal cells. These data are consistent with knockdown experiments (shown in Figure 3) which showed that suppression of these genes in EAC cells, inhibits DNA breaks and HR activity.

**Figure 5.**
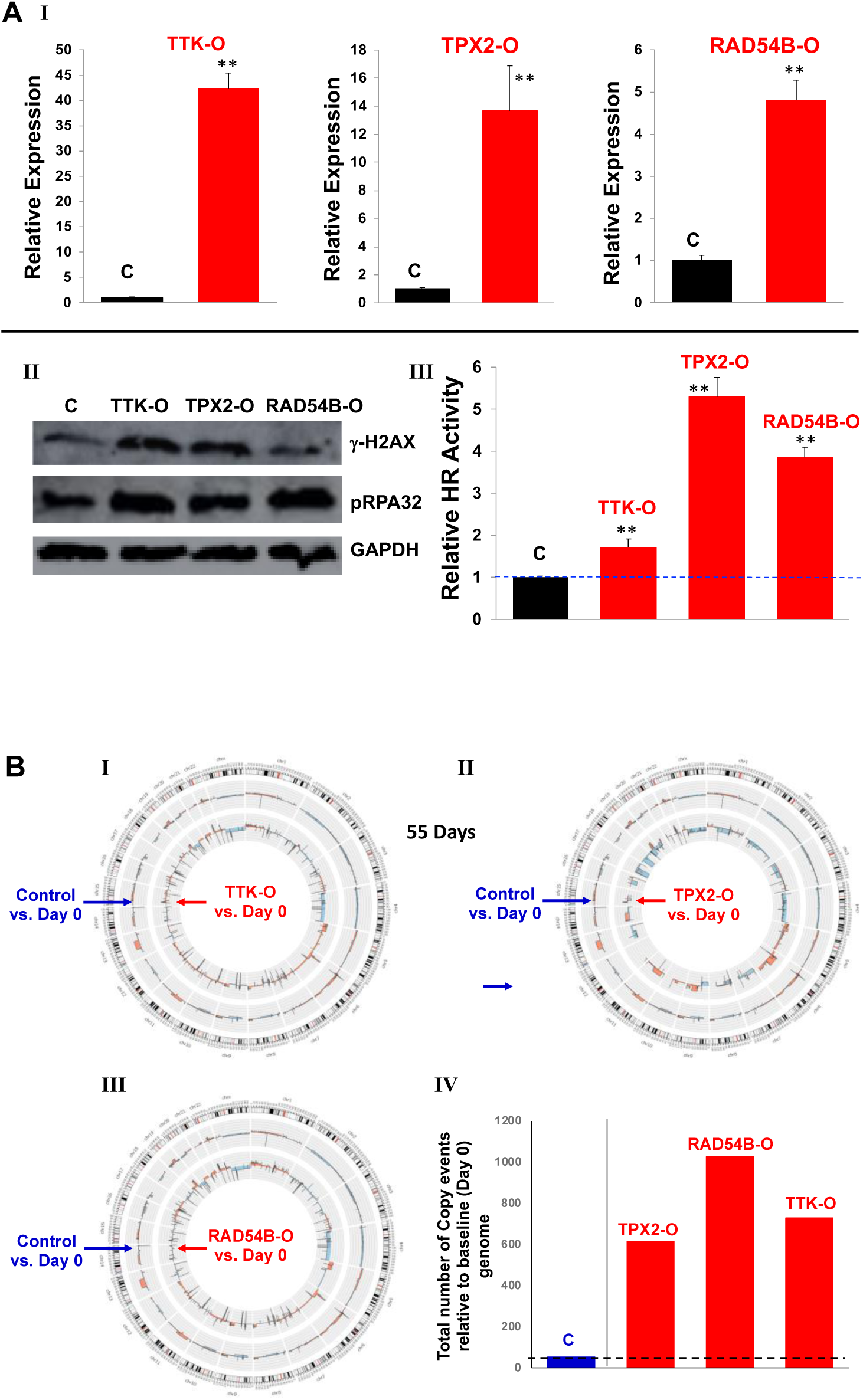
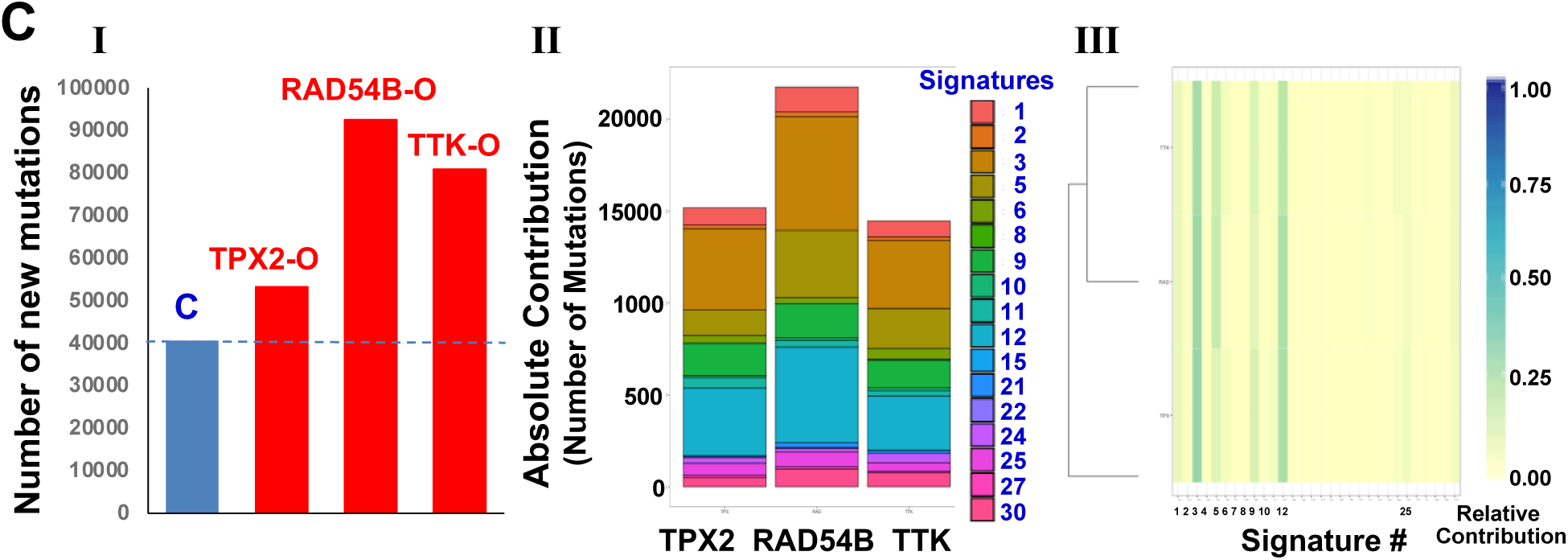
Overexpression of TTK, TPX2 and RAD54B activates mechanisms leading to genomic instability in normal esophageal cells. Normal primary human esophageal epithelial cells (HEsEpi) were transfected with control plasmid (C) or those overexpressing TTK (TTK-O), TPX2 (TPX2-O) or RAD54B (RAD54B-O), selected in puromycin and evaluated for various aspects of genome stability at indicated time points. **(A) Impact on DNA breaks and HR activity**. At day 7 after transfection, the transgene overexpression was confirmed by Q-PCR (I), and cells evaluated for **γ**-H2AX and phosphorylated-RPA32, using Western blotting (II), and HR activity, using a plasmid-based assay (III). Error bars indicate SDs of experiments conducted in triplicate. Two-tailed p values: ** = p < 0.001. **(B) Impact on genomic instability assessed using whole genome sequencing (WGS). (I-III)** Circos plots show acquisition of copy number events in control (outer circle) and transgene-overexpressing cells (inner circle) acquired in sixty days, relative to baseline genome of day 0 cells; **(IV)** Bar graph showing total copy number segments (amplifications and deletions) throughout genome in these cells. **(C) Mutational instability and underlying processes**. WGS data shown in panel B was also analyzed for mutations and mutational processes. **(I)** Total number of mutations relative to parental genome (day 0 sample); the mutations found in “day 0 sample” were removed from all samples; **(II)** Representation of mutational processes extracted. For each sample, the contribution of the 30 COSMIC signatures (https://academic.oup.com/nar/article/47/D1/D941/5146192) to the total number of nucleotide variants is shown; color assignment for each signature is shown on right; **(III)** Hierarchical clustering of the three overexpression samples according to the relative contribution of the COSMIC signatures.

### Whole genome sequencing (WGS) confirms the role of identified genes in genomic instability and provides evidence of underlying mechanism/s

Normal primary human esophageal epithelial cells transduced with control plasmid or those overexpressing TTK, TPX2 or RAD54B (shown in **Figure 5A, I**), were cultured for sixty days. DNA from these and “day 0” cells (representing baseline genome) was purified and analyzed by WGS (30X). Genome of day 0 cells was used as baseline to identify new genomic changes acquired by transduced cells over a period of sixty days. Circos plots in **Figure 5B (panels I – III)** show new copy number events in control (outer circle) and transgene-overexpressing cells (inner circle), relative to day 0 cells. The bar graph (panel IV) summarizes total copy number segments over whole genome in these cells. Relative to baseline genome (of day 0 cells), the control plasmid-transduced cells acquired 53 events in sixty days, whereas overexpression of TPX2, RAD54B and TTK led to acquisition of 614, 1026 and 729 events, respectively (Figure 5B). This shows that the overexpression of these genes caused 12 − 19-fold increase in copy number events over time, indicating a marked increase in genomic instability. Consistent with WGS data, the evaluation by SNP6.0 arrays at an earlier time point (day 30) also demonstrated that the overexpression of these genes increases genomic instability (**Supplementary Figure 3**) in normal esophageal cells.

WGS data was also analyzed for mutations. Overexpression of TPX2, RAD54B and TTK increased the mutational load in normal esophageal cells by 1.3-fold (>10,000 new mutations), 2.3-fold and 2-fold, respectively (**Figure 5C, I**). The somatic mutations in a cancer genome bear specific signatures or scar marks of distinct mutational processes or the mechanisms which give rise to these mutations. Considering all possible combinations of substitutions in a trinucleotide context, specific mutational signatures (indicative of underlying mutational processes) have been identified^21^. Using our previously published method^22,23^, we extracted the mutational signatures activated by overexpression of TPX2, RAD54B and TTK in normal esophageal cells. Although we observed contribution from at least 17 different signatures (**Figure 5C, II**), three main signatures, the Signature #3 (DNA double-strand break), #5 (of unknown aetiology) and # 9 (AID or activation-induced cytidine deaminase, a gene that induces somatic hypermutation and class-switch recombination) covered the majority of the mutational repertoire (**Figure 5C, III; Supplementary Figure 4**). This is consistent with loss and gain of function studies showing the role of these genes in increased spontaneous DNA breaks and HR activity (a repair mechanism for DNA double-strand breaks).

Thus WGS, SNP and other functional data confirmed that the overexpression of these genes caused a massive genomic instability enabling these cells to acquire a variety of genomic changes during growth in culture.

### A small molecule inhibitor of TTK inhibits EAC cell growth in vitro and in vivo, and increases efficacy of chemotherapeutic agents

Normal or non-cancerous cell types (HEsEpi, primary human esophageal epithelial cells; Het-1A, SV40 large T antigen transfected human esophageal epithelial cells; and HDF, human diploid fibroblasts) and EAC cell lines (FLO-1, OE19 and OE33) were treated with TTK inhibitor at various concentrations for 72 hr and cell viability assessed. IC50 values of the inhibitor was much higher for non-cancerous cells (HEsEpi, 252 μM; Het-1A, 5.5 μM; HDF, 3.6 μM), whereas 1.0 μM for all three EAC cell lines (**Figure 6A**). To evaluate the efficacy of TTK inhibitor in vivo, FLO-1 cells were injected subcutaneously in SCID mice and following appearance of palpable tumors, mice treated with the drug. Average tumor size in treated mice was significantly smaller than that in control mice (p = 0.01; Figure 6B).

**Figure 6:**
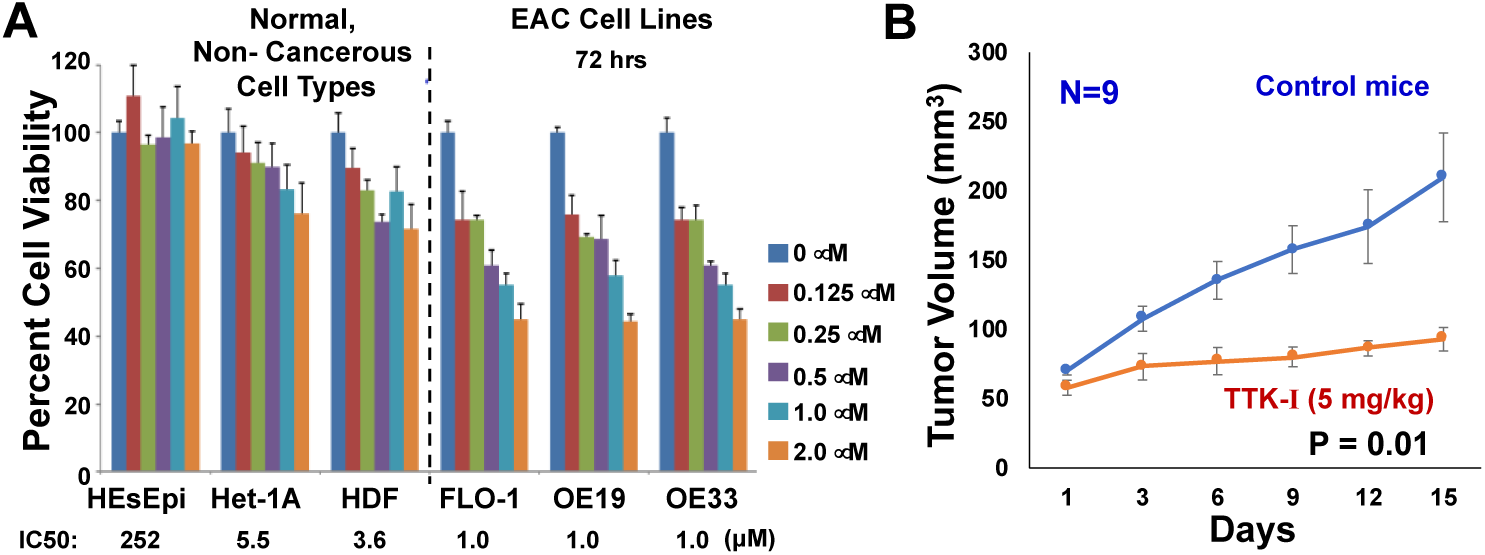
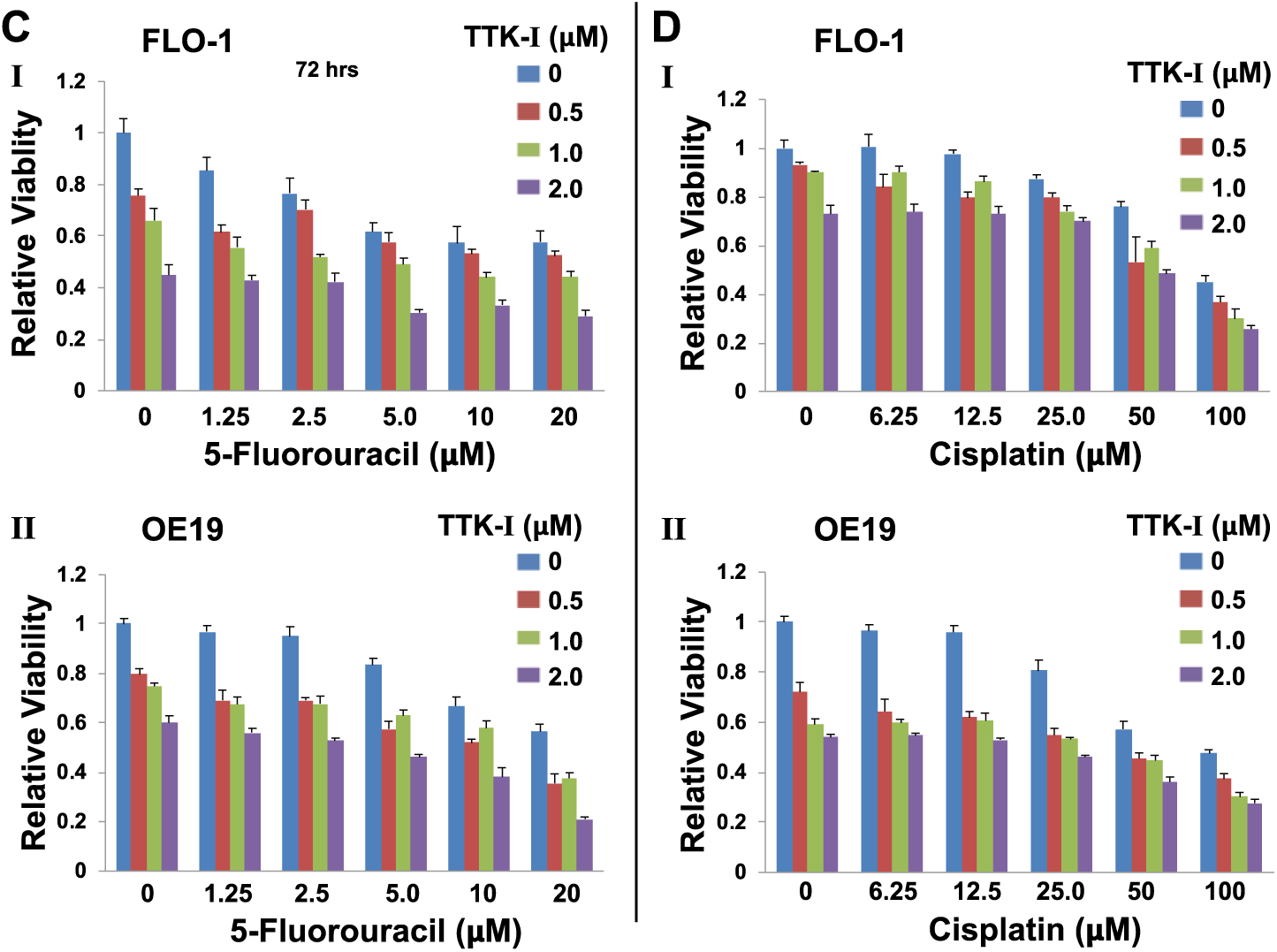
A small molecule inhibitor of TTK inhibits EAC cell growth in vitro and in vivo, and increases efficacy of chemotherapeutic agents. **(A)** Normal or non-cancerous cell types (HEsEpi, primary human esophageal epithelial cells; Het-1A, SV40 large T antigen transfected human esophageal epithelial cells; HDF, human diploid fibroblasts) and EAC cell lines (FLO-1, OE19 AND OE33) were treated with TTK inhibitor (TTK-I, CFI402257) at various concentrations for 72 hr and cell viability assessed. Error bars represent SDs of three experiments; **(B)** EAC (FLO-1) cells were injected subcutaneously in SCID mice and following appearance of palpable tumors, mice treated with TTK inhibitor. Tumor volumes were measured at indicated intervals. Error bars represent SDs of tumor sizes from nine control and nine treated mice. **(C-D)** EAC cell lines (FLO-1 and OE19) were treated with inhibitor (TTK-I, CFI402257), alone as well as in the presence of chemotherapeutic agents 5-fluorouracil (5-FU; panel C) or cisplatin (D), and cell viability measured after 72 hr. Combination indexes (calculated using CalcuSyn software) are shown in Supplementary Figure 5. Error bars represent SDs of three experiments.

To evaluate the impact of TTK inhibitor on efficacy of chemotherapeutic agents, EAC cell lines (FLO-1 and OE19) were treated with inhibitor, alone as well as in the presence of 5-fluorouracil or cisplatin, and cell viability measured after 72 hr. TTK inhibitor increased cytotoxicity of both the fluorouracil (Figure 6C) and cisplatin (Figure 6D). Combination index plots (Supplementary Figure 5) show that TTK inhibitor synergistically increased the cytotoxicity of both chemotherapeutic agents in both cell lines tested. In summary, these data show that TTK inhibitor inhibits EAC cell growth both in vitro and in vivo, and synergistically increases the efficacy of chemotherapeutic agents.

### Small molecule inhibitor of TTK inhibits spontaneous DNA damage and HR activity, and reverses genomic instability caused by chemotherapeutic agent, in EAC cells

EAC cell lines (FLO-1 and OE19) were treated with low doses of TTK inhibitor, alone or in the presence of chemotherapeutic agent, and substrate-attached (live) cells evaluated for various parameters of genome stability. TTK inhibitor reduced HR activity in EAC cell lines in a dose-dependent manner (**Figure 7A**). DNA breaks (as assessed from **γ**-H2AX expression) were reduced by TTK inhibitor, whereas increased (> 10-fold) by etoposide (**Figure 7B**). Importantly, the addition of TTK inhibitor caused a marked (5-fold) inhibition/reduction of etoposide-induced DNA breaks and DNA end resection (as assessed from pRPA32 expression) (Figure 7B). EAC cell lines treated with TTK inhibitor and/or etoposide were also evaluated for impact on micronuclei, a marker of genomic instability. In FLO1 cells, etoposide caused an increase in the micronuclei (by 12-fold), whereas a low dose of TTK inhibitor caused ∼ 3.0-fold reduction in etoposide-induced micronuclei (Figure 7C, I-II). Similarly, in OE19 cells, etoposide caused an increase in micronuclei (by 5.7-fold), whereas treatment with a low dose of TTK inhibitor not only inhibited spontaneous micronuclei (by 2-fold) but also caused 2.6-fold reduction in etoposide-induced micronuclei (Figure 7D, I-II). Taken together, these data indicate that TTK inhibitor can potentially reduce spontaneous as well as chemotherapeutic agent-induced DNA damage and genomic instability in EAC cells.

**Figure 7:**
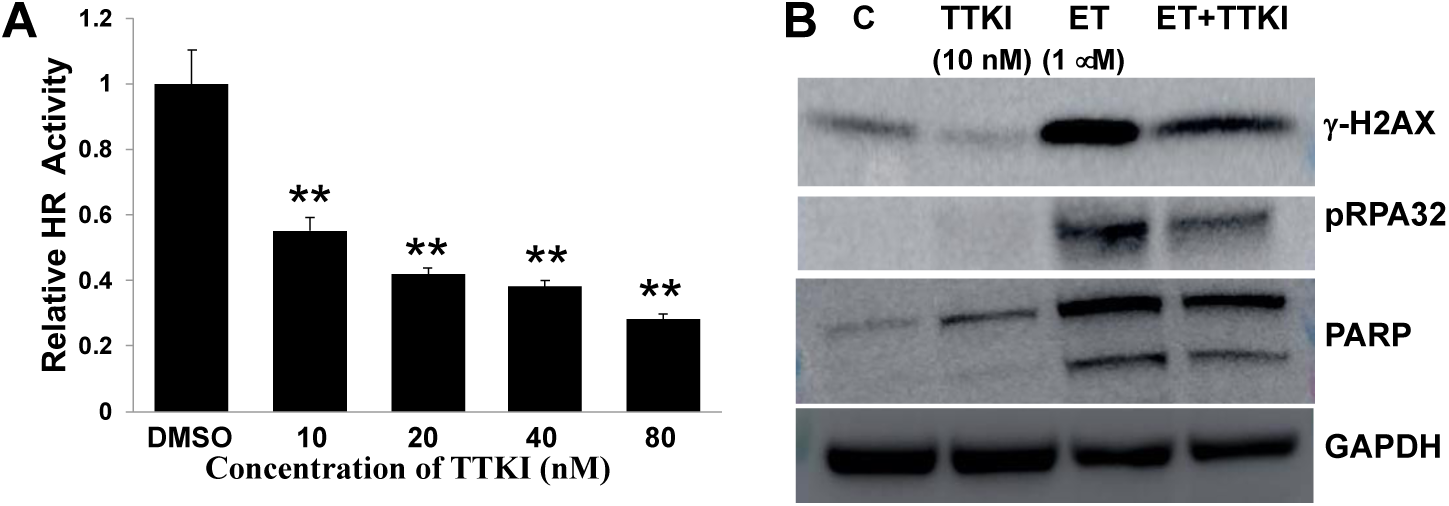
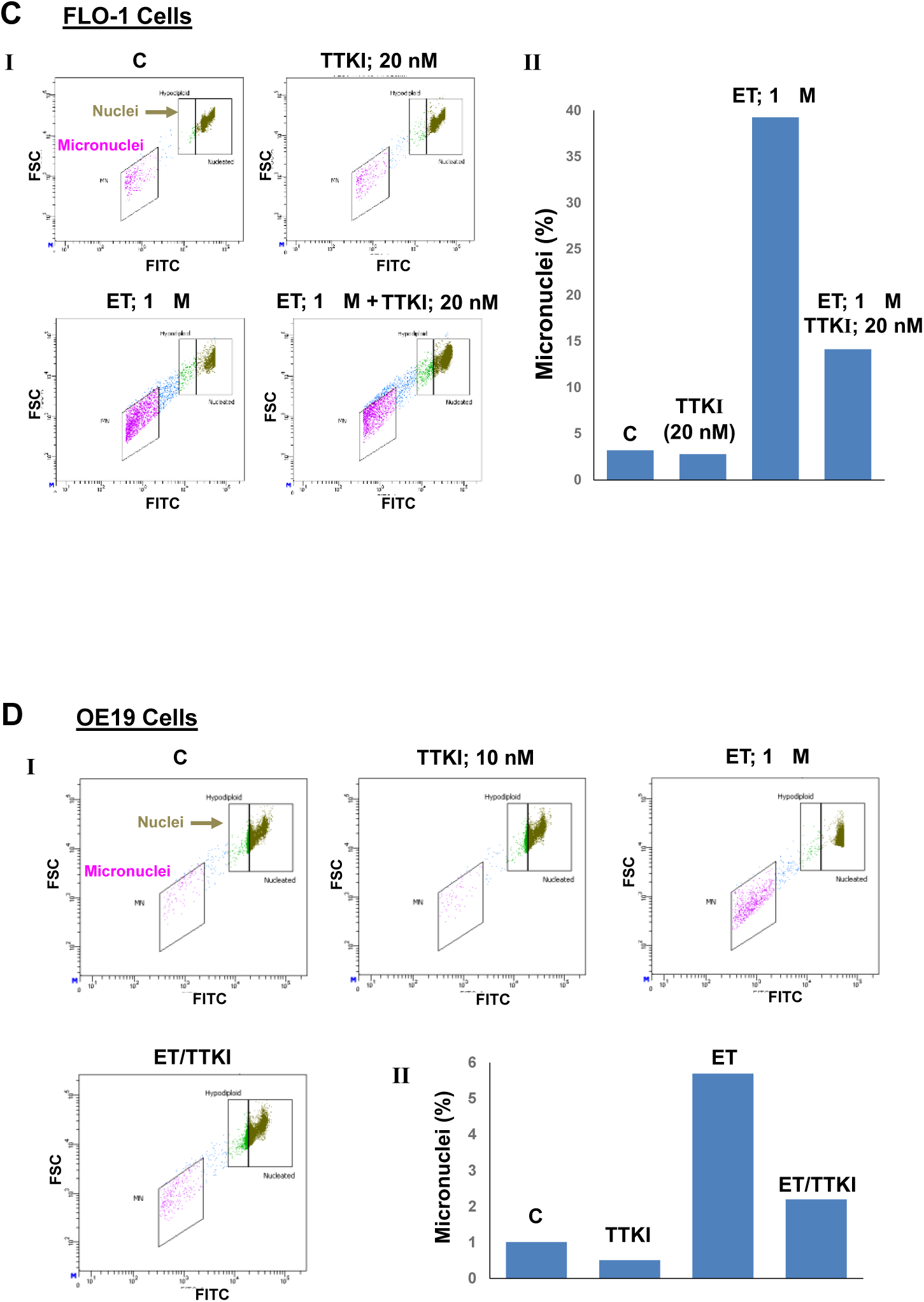
Small molecule inhibitor of TTK inhibits spontaneous HR activity and DNA breaks, and reverses genomic instability caused by chemotherapeutic agent, in EAC cells. (A) TTK inhibitor inhibits HR activity in EAC cells. FLO-1 cells were treated with different concentrations of TTK inhibitor (TTK-I, CFI402257) for 48 hr and impact on HR activity evaluated using a plasmid-based assay; Error bars indicate SDs of experiment conducted in triplicate. Two-tailed p values: ** = p < 0.001. **(B) TTK inhibitor reduces etoposide-induced DNA breaks and end resection in EAC cells**. FLO-1 cells were treated with TTK inhibitor (TTK-I) and etoposide (ET), alone as well as in combination with each other (as indicated) for 48 hr, and impact on levels of **γ**-H2AX, PARP and phosphorylated-RPA32 evaluated by Western blotting. **(C-D) TTK inhibitor reverses etoposide-induced genomic instability in EAC cells**. EAC cells, FLO1 (C) and OE19 (D), were treated with TTK inhibitor (TTK-I) and etoposide (ET), alone as well as in combination, as described in panel B and impact on micronuclei (a marker of genomic instability) assessed. Flow cytometry images of micronuclei (I) and bar graphs showing percentage of micronuclei (II) are shown.

## Discussion

Genomic instability enables a cell to constantly acquire genomic changes, some of which underlie the initiation and/or progression of cancer to advanced stages of disease. Like most other cancers, a marked genomic instability has also been observed in EAC and its premalignant states^3,4,9,10^. Consistently, the EAC genome is highly aberrant^9^ and the cancer is heterogeneous and chemoresistant^11^. Therefore, identification of genes which drive genomic evolution can help develop effective strategies for prevention and treatment of EAC.

Using EAC and multiple myeloma (MM) as model systems, we previously showed that dysregulated homologous recombination (HR) significantly contributes to genomic instability^14,16^, development of drug resistance^16^, and tumor growth^17^. Here, we used an integrated genomics approach to identify novel drivers of genomic evolution in EAC. Since genomic instability is associated with cancer progression and some of the mechanisms of genomic instability (such as HR) are also required for tumor growth, we hypothesized that if elevated expression of a gene correlates with increased genomic instability and poor survival in EAC patients, it could be a potential driver of genomic evolution. Although downregulation of a gene could also mediate genomic instability, in this study we focused only on those whose overexpression correlated with genomic instability. This is because it is relatively easy to target a gene which is overexpressed. Integration of expression, copy number and survival information (in TCGA dataset) identified 31 gene signature whose elevated expression correlated with poor survival in a second (different) EAC dataset as well as in in pancreatic and lung cancers and also in three different multiple myeloma datasets. This indicates that the signature or at least a subset of its genes are involved in genomic instability in other cancers as well. Consistent with this, loss and gain of function studies from our laboratory show that TPX2, TTK and RAD54B contribute to genomic instability in breast cancer and multiple myeloma cell lines (unpublished data). For subsequent validation, all 31 genes were first screened (using validated siRNAs) for impact on HR activity. Subsequently, fourteen of these genes were further evaluated in CRISPR/Cas9-mediated loss of function and/or overexpression screens for their role in HR and/or micronuclei (a marker of genomic instability^19^) as well as cell viability. Initial siRNA screen (evaluating all 31 genes), demonstrated that 61% of these genes had a functional role in dysregulation of HR activity in EAC cells. Overall, the siRNA, CRISPR/Cas9-knockdown and overexpression screens indicated that majority of the genes identified by integrated genomics had functional role in genomic instability and growth of EAC cells.

Three of these genes which had significant impact on the activities tested, had > 4-fold higher expression in EAC relative to normal tissues (in TCGA dataset) and represented diverse pathways (TTK, a dual specificity protein kinase; TPX2, a spindle assembly factor required for normal assembly of mitotic spindles and microtubules and RAD54B, a recombination protein), were selected for further evaluation. Suppression of all three genes reduced DNA breaks and HR activity in EAC cells. The process of HR is initiated by exonucleolytic resection resulting in the formation of a 3’-ssDNA overhang which is immediately quoted by phosphorylated-RPA32 (pRPA). Thus pRPA serves as DNA end resection marker^20,24^. Suppression of all three genes led to inhibition of DNA end resection, suggesting that these genes contribute to HR at some early step.

RAD54B is a known HR protein^25^ and its elevated expression is associated with poor prognosis in colorectal^26^ and lung^27^ cancers. Consistently, our data in EAC shows that elevated RAD54B expression contributes to increased HR activity and genomic instability, and correlates with poor survival. Role of TPX2, a microtubule associated protein, was also confirmed in HR in EAC cells. TPX2 has been shown to contribute to survival in ovarian^28^ and endometrial^29^ cancers. Cancer cells with increased genetic instability also show increased sensitivity to suppression of TPX2^30^. Our data demonstrate a novel role of TPX2 in HR in EAC. Moreover, the evaluation by both the SNP arrays as well as WGS show that overexpression of TPX2 induces genomic instability in normal esophageal cells. This is consistent with the association of TPX2 with progression in prostate cancer patients^31^. Finally, the evaluation of TTK, the spindle assembly checkpoint kinase, also confirmed its role in dysregulation of HR and genome stability in EAC. This is also consistent with a recent report demonstrating the role of TTK in HR in breast cancer^32^. TTK has been identified as one of the promising candidates for vaccination in esophageal cancer patients with advanced stage disease^33,34^. Role of TTK in survival of cancer cells and cytotoxicity of its inhibitors has also been demonstrated in hepatocellular carcinoma^35^ and glioblastoma^36^.

Although TPX2, TTK and RAD54B seem to represent diverse biological pathways, the overexpression of all three genes was associated with increased DNA damage, HR activity and genomic instability. This is also consistent with the signatures of mutational processes extracted from WGS data. Signature 3 (DNA double-strand break) and Signature 9 (AID or activation-induced cytidine deaminase, a gene that induces somatic hypermutation and class-switch recombination) were among top signatures covering majority of mutational landscape. Features of Signature 3 (https://cancer.sanger.ac.uk/cosmic/signatures_v2) (including loss of **repair** of DNA double-strand breaks by HR and increase in large deletion/insertion events which exhibit a microhomology at their breakpoint sites) suggest that it indicates aberrant/dysregulated HR. Consistently, our loss and gain of function studies also confirm that overexpression of these genes contributes to increased HR activity. Similarly, Signature 9 indicates activity of AID (activation-induced cytidine deaminase), a gene that creates mutations in DNA by deamination of cytosine to uracil and contributes to somatic hypermutation and class-switch recombination. Our unpublished data show that overexpression of AID increases DNA breaks and HR activity in normal and cancer cells. Although increased HR could be attributed to increased DNA breaks, the possibility of a functional link between AID and HR cannot be ruled out. Our functional screens, subsequent loss and gain of function studies, and previous investigations in EAC^14,15^, suggest that dysregulated HR is probably a common or predominant mechanism which drives genomic instability in EAC and possibly in other cancers^16^. Evaluation of structural variations in human genomes has identified non-allelic (aberrant) HR as one of the top underlying mechanisms^37^. Although HR is the most precise DNA repair system in normal cells, its dysregulation can lead to unnecessary and/or aberrant forms of recombination events (such as homeologous, non-allelic HR etc.) leading to genomic instability in cancer (reviewed in^38^).

Evidence from pancreatic and breast cancers suggests that the mechanism of cell death following treatment with TTK inhibitors is induction of genomic instability^35,39^. This seems to be in contradiction with our results in EAC showing that treatment with TTK inhibitor reduces spontaneous DNA damage and genomic instability. However, it should be noted that we removed dead cells and evaluated the genomic impact of TTK inhibitor in cells which survived the treatment. This is extremely important because cell death/apoptosis is also associated with DNA breaks which can hinder/confuse the results during evaluation of the impact of treatment on genomic integrity/stability. This is also the reason that we had to use lower doses of TTK inhibitor (where cell death is minimum) to accurately determine its impact on genomic integrity and stability. Treatment with TTK inhibitor at low doses inhibited both the spontaneous DNA breaks as well as genomic instability in EAC cells. This is consistent with our other observations showing that transgenic suppression of TTK in EAC cells, inhibits spontaneous DNA damage and HR activity, whereas transgenic overexpression of TTK in normal esophageal epithelial cells increases spontaneous DNA damage, HR activity and genomic instability as assessed by whole genome SNP arrays and sequencing. However, the difference between our data and that from other cancers^35,39^ could also be attributed to different mechanisms operative in these cancers. Moreover, the role of TTK in prostate cancer progression^40^ is also consistent with our data demonstrating that TTK overexpression dysregulates HR and genome stability in EAC cells.

Knockdown screen indicated that the genes identified as potential drivers of genomic evolution were also involved in EAC cell viability. Moreover, transgenic suppression of TTK, TPX2 and RAD54B not only inhibited EAC cell growth in vitro, but also significantly inhibited their growth as tumors in SCID mice. The treatment of EAC cells with a TTK inhibitor impaired their growth and synergistically increased cytotoxicity of chemotherapeutic agents. Importantly, the inhibitor of TTK inhibited spontaneous as well as chemotherapy-induced genomic instability in EAC cells. This seems to be in contrast to a general notion that genomic chaos must be enhanced (not diminished) to kill cancer cells. This is certainly true for chemotherapeutic agents as they mostly kill cancer cells by inducing DNA damage. However, this is not an ideal approach of killing cancer cells because the treatment with DNA damaging agents not only disrupts genomic integrity of normal cells but also increases DNA damage in surviving cancer cells, thus predisposing them to increased genomic instability and its harmful impact. Alternatively, the genes involved in genomic instability and growth of cancer cells (such as TTK and those identified here) can be targeted to initiate growth arrest in cancer cells without first increasing genomic instability or chaos. When such inhibitors are combined with chemotherapy, the chemotherapy-induced cytotoxicity increases while genomic instability is reduced. This is because when cancer cells (with already increased DNA breaks) are subjected to chemotherapy (such as etoposide), a large fraction of cells acquire a very high level of DNA breaks and are killed. However, there is also a fraction of cells which acquire a small or moderate increase in DNA breaks. The increase in DNA breaks in these cells is also associated with increase in HR activity, which not only helps in their survival (by reducing the number of DNA breaks) but also contributes to increase in genomic instability by (utilizing some of these breaks in) unnecessary/aberrant recombination events. When etoposide is combined with TTK inhibitor, the HR activity is reduced. This results in an increase in the fraction of cells with extensive DNA breaks (because of reduced HR), leading to increased cell death. However, the reduced HR activity in these surviving cells results in a more stable genome, as demonstrated in Figure 7 (A-D). Consistently, we have demonstrated that inhibition of HR in EAC cells, inhibits genomic instability^14^ and tumor growth^17^. Our recent data in multiple myeloma has also demonstrated that inhibition of APEX1 nuclease, a gene which drives genomic evolution in myeloma, increases cytotoxicity while inhibiting genomic instability caused by a chemotherapeutic agent^41^. We have made similar observations when HR inhibitors are used in combination with other chemotherapeutic agents (unpublished data).

## Conclusion

The integrated genomics approach used here can identify novel drivers of genomic evolution. RAD54B, TTK and TPX2 are identified as novel genes involved in dysregulation of homologous recombination and genome stability in EAC. Small molecule inhibitors of TTK and other genes identified in this study have potential to inhibit/delay genomic evolution and tumor growth. Such inhibitors also have potential to increase cytotoxicity while reducing harmful genomic impact of chemotherapy.

## Materials and Methods

### Patient dataset

EAC patients (n=88) from The Cancer Genome Atlas data.

### Integrated genomics

We hypothesized that if elevated expression of a gene correlates with increased genomic instability and poor survival of a patient, it could be a potential driver of genomic evolution. To identify these genes, we used following stepwise process; details of each step are provided in supplementary material: **1)** Investigated gene expression in 11 normal and 88 EAC patient samples in TCGA dataset and identified genes that were overexpressed in EAC; **2)** Assessed genomic instability in patient samples by counting total number of copy events in each patient. Integrated genomic instability data with expression data to identify genes that were overexpressed in EAC and whose expression correlated with genomic instability; **3)** Evaluated resulting list of genes for correlation with survival. This led to identification of 31 genes that were overexpressed in EAC and whose expression correlated with genomic instability and overall survival in EAC patients in TCGA dataset; **4)** The expression of this 31 gene signature was tested for correlation with survival in a second EAC dataset as well as in other cancers.

### Functional screens to validate genomic instability signature

Expression of thirty one genes, identified as potential drivers of genomic instability in EAC (Figure 1) was either suppressed (using validated siRNAs as well as CRISPR/Cas9 technology) or overexpressed (using open reading frame expression plasmids in viral constructs) in EAC (FLO-1) cells, and impact on micronuclei (marker of genomic instability), homologous recombination (HR; a mechanism of genomic instability in EAC) and cell viability, assessed.

### Constructs for loss and gain of function and transductions/transfections

Validated siRNAs and lentiviral constructs expressing shRNAs were purchased from Sigma. Constructs for CRISPR/Cas9-mediated gene knockout (Cas9 and guide RNAs) and plasmids expressing open reading frames (for overexpression) were obtained from Dr. David Root at Broad Institute, Boston, MA. For CRISPR/Cas9-mediated gene knockout, EAC cells were stably transduced with Cas9 and then transduced with guide RNAs targeting individual genes. For shRNA-mediated gene suppression/overexpression, lentiviral constructs were transduced in normal and EAC cells as reported previously^14,15,17^.

### Cell types

Normal primary human esophageal epithelial (HEsEpiC) cells were purchased from ScienCell Research Laboratories (Carlsbad, CA); SV40 large T antigen transfected human esophageal epithelial cells (Het-1A; CRL-2692) and normal human diploid fibroblasts purchased from American Type Culture Collection (Manassas, VA); and EAC cell lines (FLO-1, OE19 and OE33) purchased from Sigma Aldrich Corporation (Saint Louis, MO). Cells were cultured as reported previously^14,15,17,42^.

### Antibodies and reagents

Anti-RPA32 (phospho Ser4, phospho Ser8) antibody was purchased from Novus Biologicals LLC (Centennial, CO) and anti-phospho-histone H2A.X (Ser139) (20E3) antibody from Cell Signaling Technology Inc., (Danvers, MA). TTK inhibitor “CFI-402257” was purchased from MedChemExpress LLC (Monmouth Junction, NJ). For evaluation of TTK, TPX2 and RAD54 expression by real time PCR, TaqMan probes from Applied Biosystems INC. (Beverly, MA) were used.

### Cell Viability

Assessed using Cell Titer-Glo Luminescent Viability Assay kit (Promega Corporation, Madison, WI).

### Evaluating impact on spontaneous DNA breaks and DNA end resection

Impact of transgenic modulations on DNA integrity was monitored by evaluating cells for **γ**-H2AX, a marker for DNA breaks, using Western blotting. Impact on DNA end resection, a critical step in the initiation of HR, was monitored by evaluating phosphorylation (on ser4 and ser8) of RPA32^20^.

### Homologous recombination (HR) activity

HR activity was measured in a plasmid substrate, using a luminescence-based assay reported by us previously^14,17,41^.

### Evaluating impact on genomic evolution

#### Micronucleus assay

Micronuclei (marker for genomic instability^19^) were evaluated by flow cytometry, using a commercial kit as reported by us previously^41^.

#### *Whole genome sequencing (WGS) and* single nucleotide polymorphism (SNP) arrays

Briefly, the control and transgenically-modulated cells were cultured for different durations, genomic DNA purified and analyzed using SNP6.0 arrays (Affymetrix) or WGS platforms. For each experiment, the parental or “day 0” cells (saved at the beginning of experiment) were used as baseline genome to detect changes in control and transgenically-modulated cells, during their growth in culture. SNP and WGS data were analyzed as described by us previously^14,16,22,23^.

### Evaluating cell growth in a subcutaneous tumor model

Six to twelve week-old CB17/ICr-SCID mice were maintained as per guidelines of the Institutional Animal Care and Use Committee (IACUC). All experimental protocols were reviewed and approved by the IACUC and the Occupational Health and Safety Department of Dana Farber Cancer Institute, Boston, MA. The mice were exposed to 150 rads x-irradiation and injected subcutaneously with 5×10^6^ EAC (FLO1) cells, either untransduced (for evaluation of TTK inhibitor) or transduced with shRNAs (control or those targeting TPX2, TTK and RAD54). Transduced cells were selected in puromycin and cultured for another two weeks prior to inoculation in mice. For evaluation of TTK inhibitor, following appearance of palpable tumors, mice were treated with vehicle control or inhibitor (5 mg/kg injected intraperitoneally five times a week for three weeks). Tumor sizes were measured at least three times every week.

## Acknowledgements/Funding

This work was supported by Department of Veterans Affairs Merit Review Award I01BX001584-01 (NCM), NIH grants P01-155258 and 5P50 CA100707 (MAS, MKS, NCM) and Leukemia and Lymphoma Society translational research grant (NCM).

## Author contributions

MAS envisioned the study, analyzed and interpreted data and prepared manuscript; NCM assisted in data interpretation and critical review of manuscript; LB, MR and MKS conducted bioinformatic and statistical analyses; SK, LB and ST equally contributed to major experiments and manuscript preparation; CL, JS, CC, GBG and RP contributed to specific experiments and data analyses.

