## Supplementary Material for "Integrated genomics and comprehensive validation reveal novel drivers of genomic evolution in esophageal adenocarcinoma"

### METHODS:

#### Details of each step in the identification of genomic instability signature:

**Identification of differentially expressed genes:** RNASeq raw counts of TCGA data (containing 95 cases for esophageal squamous cell carcinomas, 88 cases of EAC and 11 normal samples: Level 3, Illumina Hiseq RNASeqV2) were downloaded and pre-processed separately, using the package TCGAbiolinks[1]. Each subtype expression matrix was normalized using the TCGAanalyze_Normalization function encompassing the EDASeq protocol[2]. Differential expression was computed with the TCGAanalyze_DEA function, implementing the EdgeR protocol[3]. **Overexpressed genes:** A cutoff of 1 was set for log2 fold change (in EAC relative to average of normal samples) with an FDR of 0.01 to identify genes overexpressed in EAC. Only EAC data was subsequently used in this study.

**Evaluation of copy number events in each patient:** Genomic instability in each patient was assessed from total number of copy number events, which were computed using an R script. A copy event was defined as “high” if the segment mean was $\geq$ 2.5 and “low” if the segment mean was $\leq$ 1.5.

**Survival Analysis:** Overall survival was defined as the day of last follow-up or day of death depending on vital status. Samples with a value of “NA” in the vital status row were excluded. Univariate analysis was performed using Cox regression with gene expression as the independent variable. For each gene, we recorded regression coefficient and hazard ratio with P-value cutoff of 0.05.

### RESULTS:

**Evaluation by single nucleotide polymorphism arrays confirms the role of identified genes in genomic instability.** Normal primary human esophageal epithelial cells transduced with control plasmid or those overexpressing TTK, TPX2 or RAD54B (shown in Figure 4A, I), were cultured for thirty days. DNA from these and “day 0” cells (representing baseline genome) was purified and analyzed by single nucleotide polymorphism (SNP6.0) arrays (Affymetrix). Genome of “day 0” cells was used as baseline to identify new genomic changes acquired by transduced cells over a period of thirty days. Relative to baseline genome, the overexpression of all three genes was associated with the acquisition of new copy number events throughout chromosomes (Supplementary Figure 3A), ranging from 4.4- to 4.9-fold increase relative to control shRNA-transduced cells (Supplementary Figure 3B). Thus, evaluation by WGS (at day 60) as well as evaluation by SNP arrays (at day 30) demonstrated that overexpression of these genes increases genomic instability in normal esophageal cells.

**Patterns of single nucleotide variation subdivided according to the preceding and following nucleotide.** Whole genome sequencing data shown in Figure 5 was analyzed for mutational changes as described previously[4-8]. Although we observed contribution from at least 17 different signatures (Figure 5C, II), three main signatures, the Signature #3 (DNA double-strand break), #5 (of unknown aetiology) and # 9 (AID or activation-induced cytidine deaminase, a gene that induces somatic hypermutation and class-switch recombination) covered the majority of the mutational repertoire (Figure 5C, III; Supplementary Figure 4).

**SUPPLEMENTARY FIGURE LEGENDS:**

**Supplementary Figure 1. Co-upregulated GIS31 genes in EAC.** GIS31 genes in EAC patient dataset (from TCGA) were analyzed by a pairwise correlation of their expression, using Pearson method.

**Supplementary Figure 2. Functional siRNA screen evaluating GIS31 genes for impact on homologous recombination (HR) activity in EAC cells.** EAC (FLO1) cells were transfected with siRNAs, either control (non-targeting) or those targeting 31 potential genomic instability (GIS31) genes, and impact on HR assessed using strand exchange assay described in Methods section. Bar graphs show percent inhibition of HR activity; error bars represent SDs of three independent experiments. Two-tailed p-values, indicating significance of difference relative to control siRNA-transfected cells, are shown as: * < 0.05 – > 0.005; ** < 0.005 – > 0.0001; *** < 0.0001 – < 0.000005.

**Supplementary Figure 3. Overexpression of TTK, TPX2 and RAD54B in normal primary esophageal cells induces the acquisition of new copy number changes over time.** Normal primary human esophageal epithelial cells (HEsEpi; ScienCell) were transfected with control plasmid (C) or those overexpressing TTK (TTK-O), TPX2 (TPX2-O) or RAD54B (RAD54B-O), selected in puromycin and cultured for thirty days. DNA from these and baseline control (day 0) cells was purified and acquisition of copy number events during growth of cells in culture vs. day 0 cells (representing baseline genome) monitored, using SNP6.0 arrays (Affymetrix); a copy event was defined as a change in ≥ 5 consecutive CNV probes by 1 copy. **(A)** Images showing copy number events, as red (amplification) and blue (deletion) dots on all chromosomes, in cultured relative to baseline (day 0) cells; **(B)** Bar graph showing total copy-number change events, throughout genome.

**Supplementary Figure 4. Patterns of single nucleotide variation subdivided according to the preceding and following nucleotide.** Whole genome sequencing data shown in Figure 3 was analyzed for mutational changes. **(A)** Histogram representing the fraction of mutations found in any of the 96 trinucleotide context. **(B)** Mutation model best fitting the sample SNVs based on 30 COSMIC (https://academic.oup.com/nar/article/47/D1/D941/5146192) genomic signatures; most important signatures and their contribution to the model are reported as title. (C) Mutational pattern not explained by the best fitting model based on COSMIC signatures.

**Supplementary Figure 5: A small molecule inhibitor of TTK synergistically increases the efficacy of chemotherapeutic agents.** EAC cell lines, FLO-1 and OE19 were treated with inhibitor (TTK-I, CFI402257), alone as well as in the presence of chemotherapeutic agents 5-fluorouracil (5-FU; panel I) or cisplatin (II), and cell viability measured after 72 hr. Combination indexes, calculated using CalcuSyn software, are shown.

Supplementary Figure 1

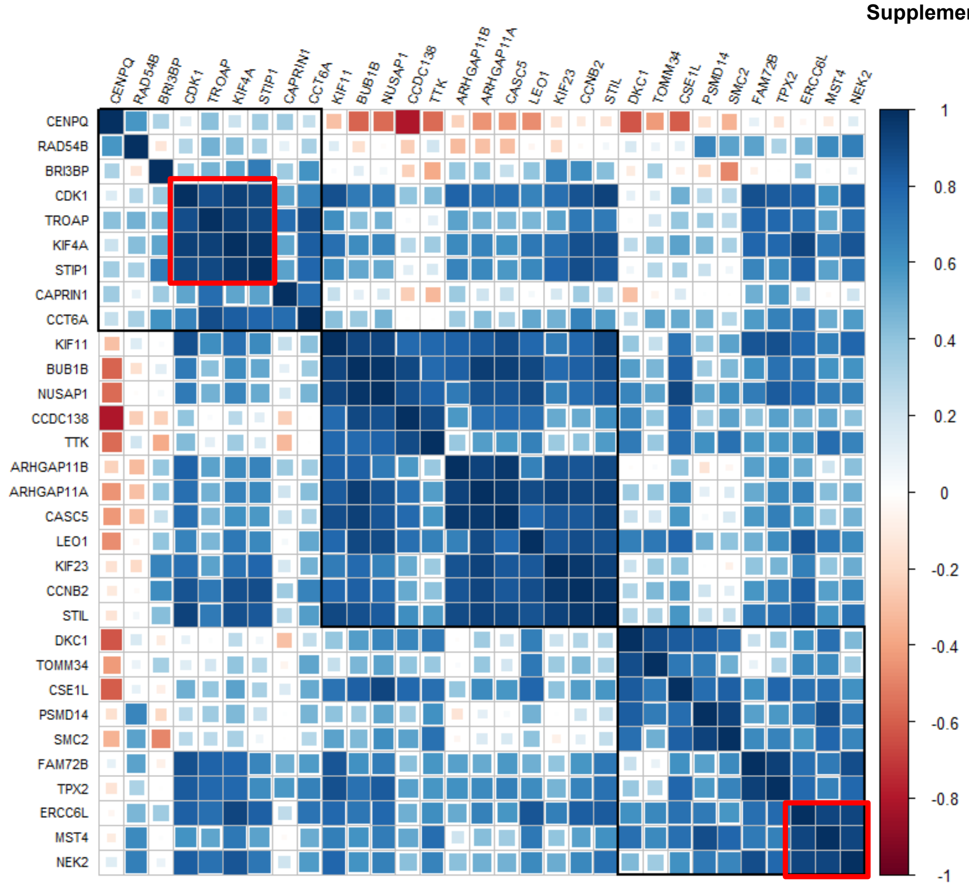

Supplementary Figure 2

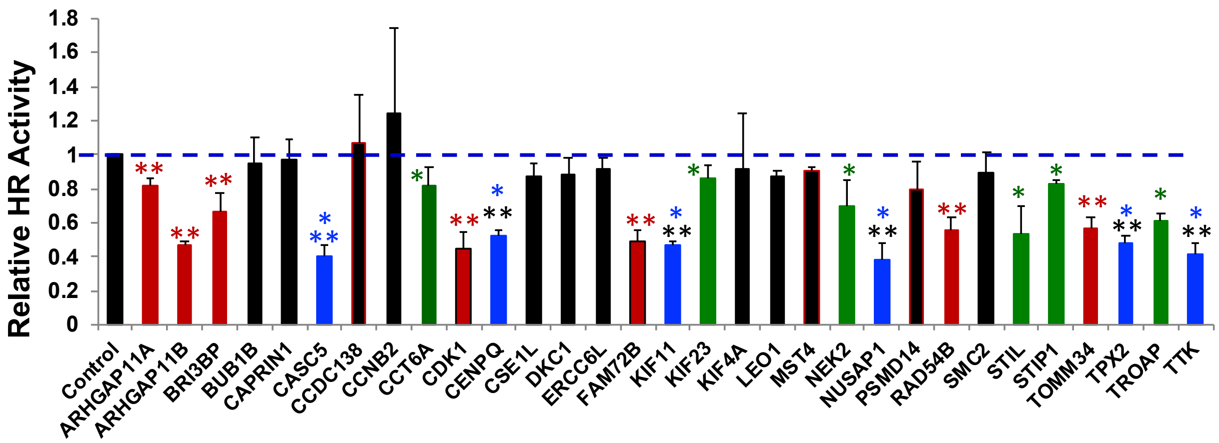

Supplementary Figure 3

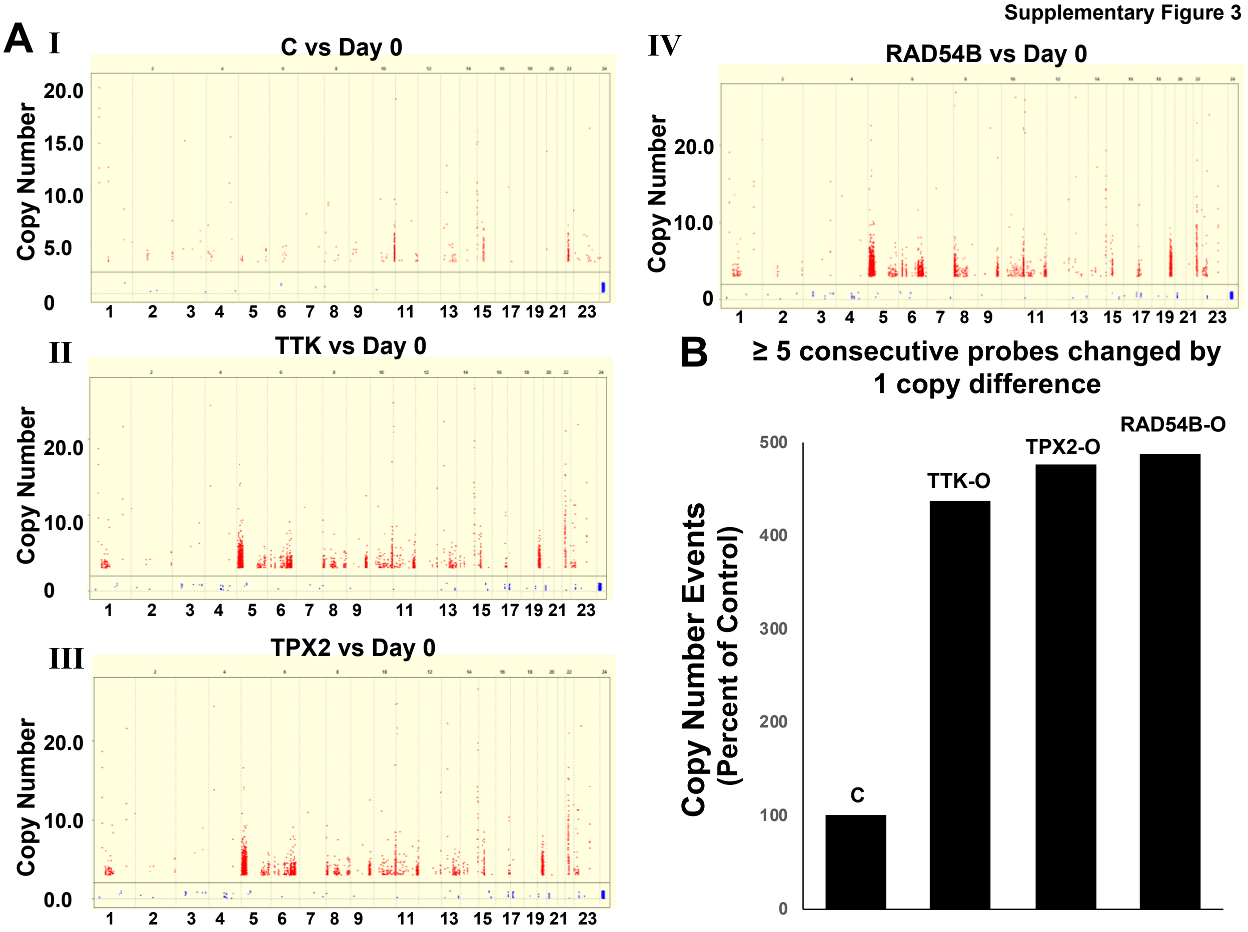


Supplementary Figure 4

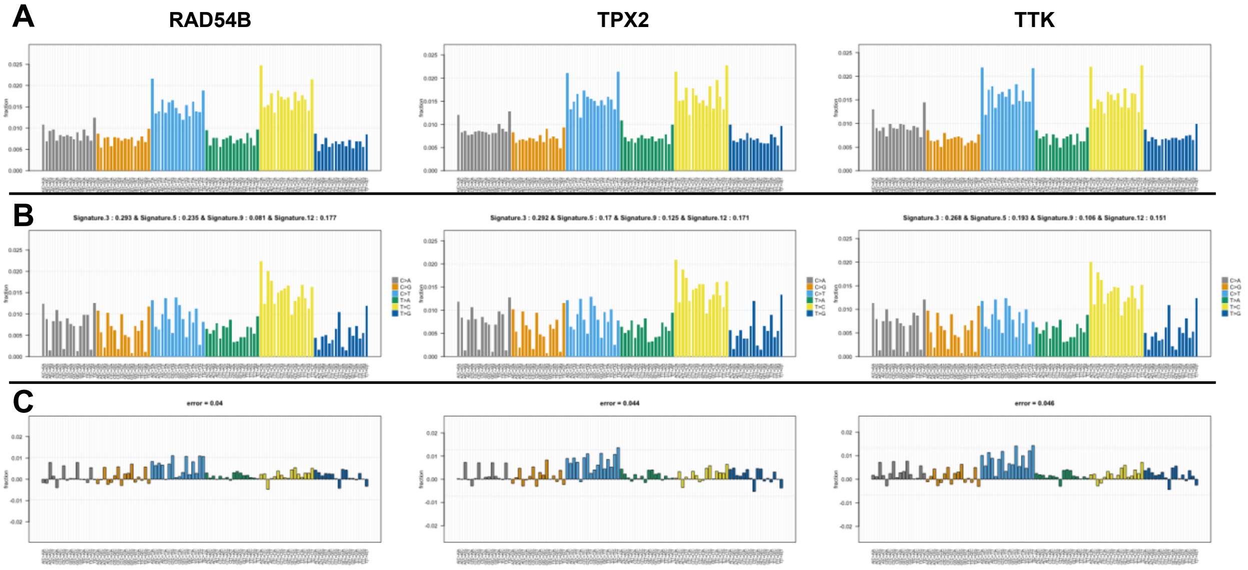


Signature 3: dsDNA break, Signature 5: unknown,
Signature 9: AID, Signature 12: unknown;

Supplementary Figure 5

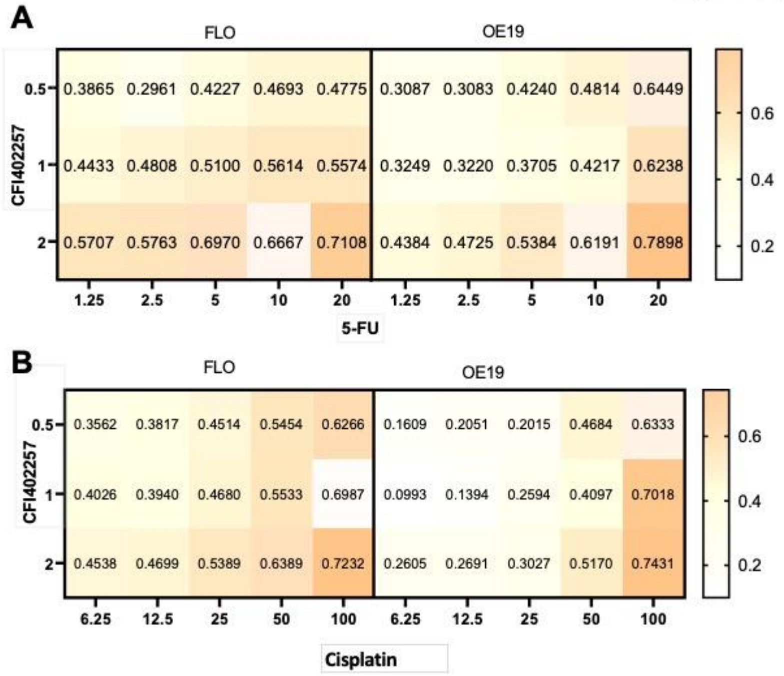
